# Identification of a Novel miRNA Expression Signature for Lung Adenocarcinoma Using Systematic Machine Learning Optimization

**DOI:** 10.64898/2026.01.10.698764

**Authors:** Shruti Agrawal, Pralay Mitra

**Author notes:** WWW home page: https://sites.google.com/view/shruti07.

## Abstract

Lung adenocarcinoma (LUAD), the most common lung cancer subtype, urgently requires reliable microRNA (miRNA) biomarkers for early detection and therapy. This study introduces a machine learning framework integrating feature stability analysis, precision-recall curves, and resampling strategies (e.g., SMOTE) to robustly identify miRNA signatures from imbalanced TCGA-LUAD data (564 samples: 519 tumor, 45 normal). We selected 8 stable features (hsa-mir-143, hsamir-210, hsa-mir-21, hsa-mir-183, hsa-mir-96, hsa-mir-182, hsa-mir-130b, hsa-mir-141) with 100% cross-fold stability via 10-fold cross-validation. A Random Forest classifier yielded excellent training performance (AUC: 1.0000; accuracy: 98%) and good generalization on an independent test set (AUC: 0.8438; accuracy: 75%). Consistent feature importance across folds supports biological relevance over overfitting. The framework mitigates class imbalance, high dimensionality, and distribution shifts—key hurdles in biomarker discovery. These reproducible miRNAs hold promise as non-invasive diagnostic tools, though external validation underscores generalization challenges across cohorts.

## 1 Introduction

Lung cancer remains the leading cause of cancer mortality worldwide, claiming 1.8 million lives annually [7, 18]. Despite advances, the 5-year survival rate lingers at 16.8%, primarily due to late diagnosis [14, 13]. While low-dose CT (LDCT) screening aids early detection, its poor specificity for benign vs malignant nodules underscores the need for molecular biomarkers [10, 18].

Lung adenocarcinoma (LUAD), comprising 40% of lung cancers and a major non-small cell lung cancer (NSCLC) subtype, displays unique molecular profiles ripe for biomarker exploitation [9]. MicroRNAs (miRNAs), stable, 21-25 nt non-coding RNAs that post-transcriptionally regulate 2,654 human mature sequences (miRBase v22), are dysregulated in LUAD, acting as oncogenes or suppressors [11, 7, 21]. Their circulation in biofluids enables non-invasive liquid biopsies for early detection, staging, and metastasis prediction [2, 15, 4].

High-throughput sequencing has generated vast multi-omics datasets like TCGA, profiling thousands of LUAD tumors. Yet, high dimensionality (thousands of miRNAs across 500 samples) fuels overfitting and poor generalization in traditional analyses. Machine learning (ML) excels here, capturing non-linear patterns via automated feature selection [6]. Supervised models like Random Forest, SVM, Gradient Boosting, and regularized regression outperform in genomic classification, yielding minimal signatures for tumor detection and prognostication [14, 10]. However, most studies overlook end-to-end pipeline optimization for miRNA-LUAD tasks, neglecting class imbalance (e.g., 11:1 tumor:normal ratios), overfitting validation, and screening-vs-diagnostic distinctions [18].

This study introduces an enhanced computational framework tailored to miRNA biomarker discovery in LUAD, addressing class imbalance, feature instability, and distributional shifts. The approach employs 10-fold cross-validation with feature selection frequency and importance tracking, imbalance-aware metrics (Precision-Recall curves and Average Precision alongside ROC-AUC), and hybrid resampling via SMOTEENN. Four classifiers, Logistic Regression, Random Forest, Support Vector Machine, and Gradient Boosting, underwent unified preprocessing and internal/external validation to evaluate generalizability. Applied to TCGA-LUAD data (imbalance ratio 11.53:1), the framework identified eight miRNAs with 100% cross-fold stability and outstanding internal performance (AUC = 1.000), while revealing external validation challenges. The full workflow is depicted in Figure 1.

**Fig. 1.**
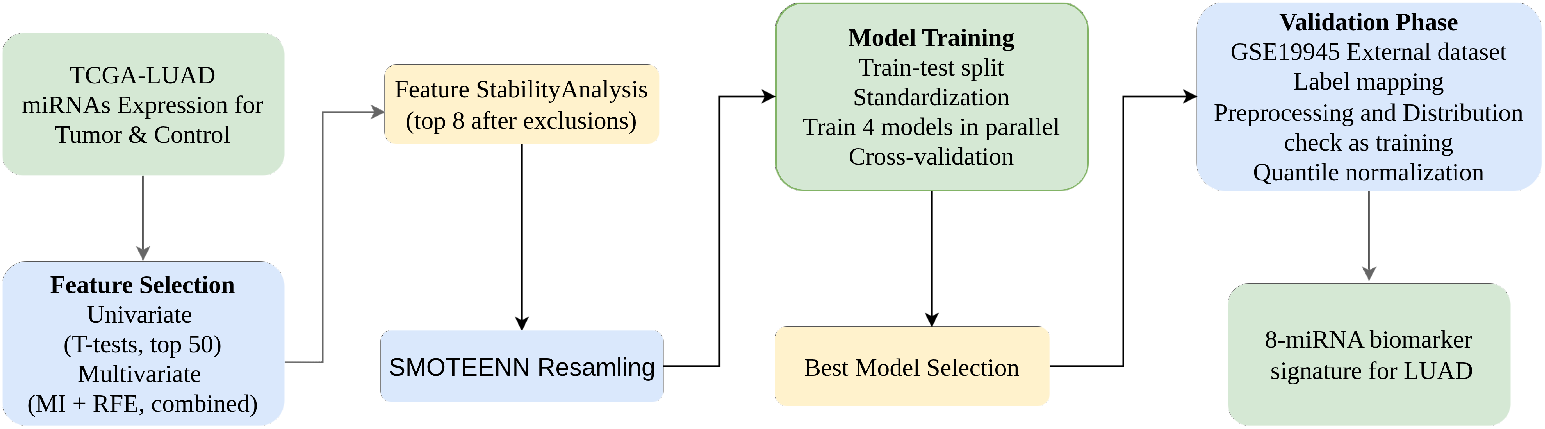
Complete stability-driven miRNA biomarker discovery pipeline for LUAD. From TCGA-LUAD (564 samples, 1881 miRNAs) through hierarchical feature selection (62 candidates), 10-fold stability analysis (8 miRNAs, S=1.0), SMOTEENN resampling, Random Forest optimization, to external validation on GSE19945 (AUC=0.8438 despite batch effects).

## 2 Methods

### 2.1 Dataset and Preprocessing

We analyzed miRNA-seq profiles from TCGA-LUAD tissue samples (564 total: 519 tumor [92.0%], 45 normal [8.0%]; 1,881 features; imbalance ratio 11.53:1). This mirrors clinical scarcity of normals but risks majority-class bias. An independent test set from GSE19945 (12 samples: 8 normal, 4 cancer; inverse distribution) assessed distributional shift robustness. Normalized expression levels were independent variables; binary status (cancer/normal) was the target.

Data preprocessing ensured robustness and downstream compatibility by verifying data integrity, imputing missing values with the median, setting negative expression values to zero, and clipping extreme outliers (*>*5 SD). A log_2_(*x* + 1) transformation stabilized variance and reduced skewness, followed by feature standardization (zero mean, unit variance) using StandardScaler for stable model training.

### 2.2 Feature Selection Strategy

We implemented a hierarchical two-stage approach:

Stage 1 – Univariate Testing: Independent t-tests with FDR correction, foldchange (cancer/normal mean), and p-value ranking selected the top 50 differentially expressed miRNAs.

Stage 2 – Multivariate Selection: Mutual information (MI; top 30 features) captured non-linear dependencies, while recursive feature elimination (RFE) with Random Forest selected the top 30 optimizing collective performance. Their union formed the robust candidate set.

### 2.3 Feature Stability Analysis

Feature stability was assessed using 10-fold cross-validation on the candidate miRNA set. Robustness was quantified using three complementary metrics.

The stability score (*S*) was defined as the proportion of cross-validation folds in which a given feature was selected:

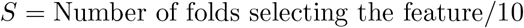

Based on this score, features were categorized as: Highly stable : *S* ≥ 0.80 ( ≥ 8 folds), Moderately stable : 0.60 ≤ *S <* 0.80 (6–7 folds), Unstable : *S <* 0.60 (*<* 6 folds).

The coefficient of variation (CV) was used to assess the consistency of feature importance across folds and was calculated as:

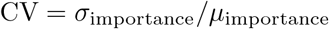

where *σ*_importance_ and *µ*_importance_ denote the standard deviation and mean of Random Forest feature importance scores, respectively. Lower CV values indicate greater robustness.

Finally, mean importance was computed as the average Random Forest importance score across all folds:

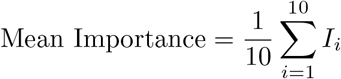

where *I*_*i*_ represents the importance score in fold *i*.

From the top 30 candidate miRNAs (Figure 2), eight miRNAs (hsa-miR-143, hsa-miR-210, hsa-miR-21, hsa-miR-183, hsa-miR-96, hsa-miR-182, hsa-miR-130b, and hsa-miR-141) achieved perfect stability (*S* = 1.0), exhibited low CV values and high mean importance, and were available in the external validation dataset (GSE19945). These features were therefore selected as the final biomarker panel, balancing predictive performance, robustness, and clinical applicability.

**Fig. 2.**
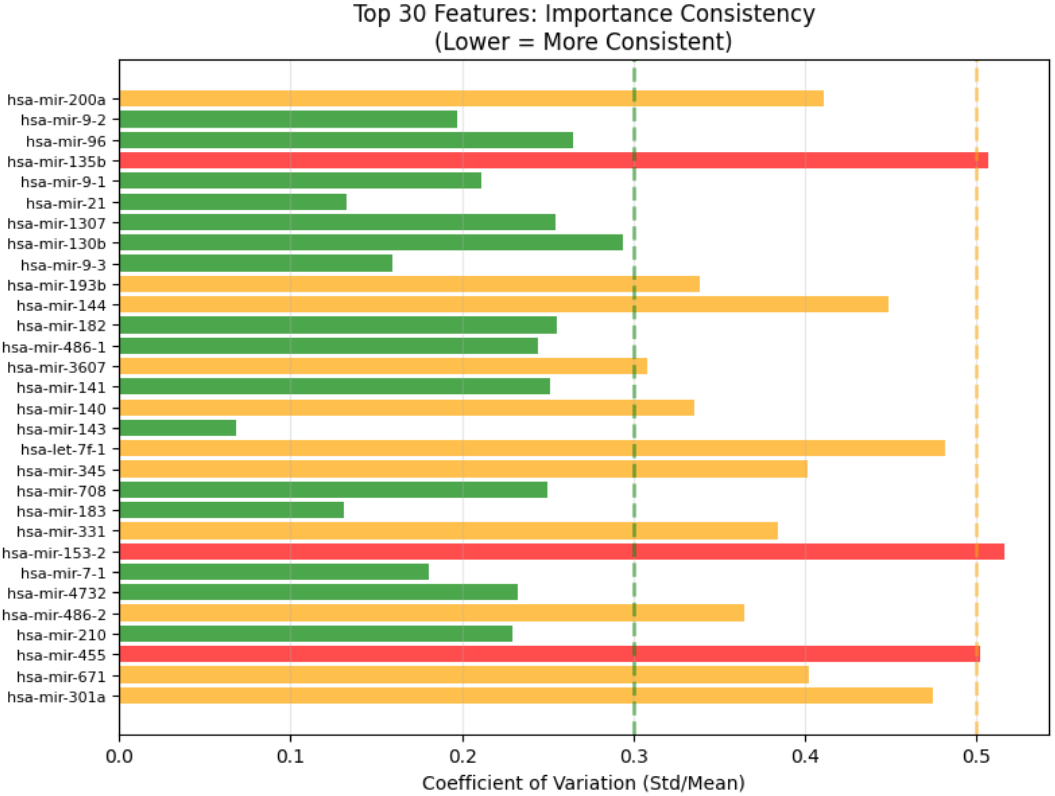
Feature importance consistency across 10-fold cross-validation. Coefficient of variation (CV = *σ*/*µ* of Random Forest importance scores) for top 30 candidate miRNAs from hierarchical selection. The final 8-miRNA panel green bars demonstrate exceptional consistency (lowest CV values), confirming robustness to data partitioning. Remaining candidates show higher variability (yellow/orange), validating stability-based selection criteria (CV *<* 0.15, S = 1.0). Lower CV indicates greater reproducibility across folds, supporting biological relevance over overfitting artifacts.

### 2.4 Addressing Class Imbalance: SMOTEENN

The training set exhibited severe imbalance (363 cancer, 31 normal; ratio 11.7:1). To address this, a hybrid SMOTEENN strategy was applied, where SMOTE (k = 5) generated synthetic normal samples via k-nearest neighbor interpolation, followed by ENN (k = 3) to remove noisy samples misclassified by the majority of nearest neighbors.

This transformed 394 samples into 710 balanced samples (349 cancer, 361 normal; ratio 1.03 : 1), an 80% increase preserving decision boundaries. The slight normal predominance was intentionally compensated for GSE19945’s inverse distribution (8 normal, 4 cancer).

### 2.5 Machine Learning Model Development

Four widely used biomedical classifiers—Random Forest (200 trees), Gradient Boosting (100 estimators), SVM with RBF kernel, and L2-regularized Logistic Regression—were evaluated on the SMOTEENN-balanced dataset using a 70:30 stratified split. Model robustness was assessed via 5-fold stratified crossvalidation, with AUC-ROC as the primary metric alongside accuracy, precision, recall, F1-score, and PR-AUC (Figure 3). Models were selected based on test AUC/AP, cross-validation stability, and balanced sensitivity–specificity. Random Forest emerged as optimal, achieving a perfect test AUC (1.000) with highly stable CV performance (0.9997 *±* 0.0006). Performance details shown in Figure 3, Table 2.

**Table 1.** Biological Relevance of the 8 Stability-Selected miRNA Biomarkers (S=1.0)

| miRNA | Chromosome | Role in LUAD | Diagnostic/Prognostic Utility |
| --- | --- | --- | --- |
| hsa-miR-143 | 5q32 | Tumor suppressor (↓)<br>targets KRAS, ERK5 | Low expression predicts poor survival [16] |
| hsa-miR-210 | 11p15.5 | HypoxamiR (↑)<br>targets ISCU, COX10 | High levels indicate hypoxia/prognosis [3] |
| hsa-miR-21 | 17q23.2 | OncomiR (↑)<br>targets PTEN, PDCD4 | Circulating biomarker, early detection [1] |
| hsa-miR-183 | 7q32.2 | OncomiR (↑)<br>targets EZR, DKK family | Elevated in advanced stages [5] |
| hsa-miR-96 | 7q32.2 | OncomiR (↑)<br>targets FOXO1, RECK | Metastasis association[8] |
| hsa-miR-182 | 7q32.2 | OncomiR (↑)<br>targets FOXO3, PTEN | Lymph node prognostic [20] |
| hsa-miR-130b | 22q11.1 | OncomiR (↑)<br>Targets: PPARγ | Serum recurrence biomarker[12] |
| hsa-miR-141 | 12p13.31 | Tumor suppressor (↓)<br>targets ZEB1/2 | Circulating metastasis predictor [17] |

**Table 2.** Performance Comparison of Four Machine Learning Classifiers on 8-miRNA Panel (Post-SMOTEENN Balance)

| Model | Test AUC | CV AUC | Accuracy | Precision | Recall | F1-score |
| --- | --- | --- | --- | --- | --- | --- |
| Logistic Regression | 0.9986 | 0.9998 | 0.98 | 0.91 | 0.99 | 0.95 |
| Random Forest | **1.000** | 0.9997 | 0.98 | 0.89 | 0.99 | 0.93 |
| Support Vector Machine | 0.9954 | 0.9998 | 0.98 | 0.91 | 0.99 | 0.95 |
| Gradient Boosting | 0.9872 | 0.9985 | 0.98 | 0.89 | 0.99 | 0.93 |

**Fig. 3.**
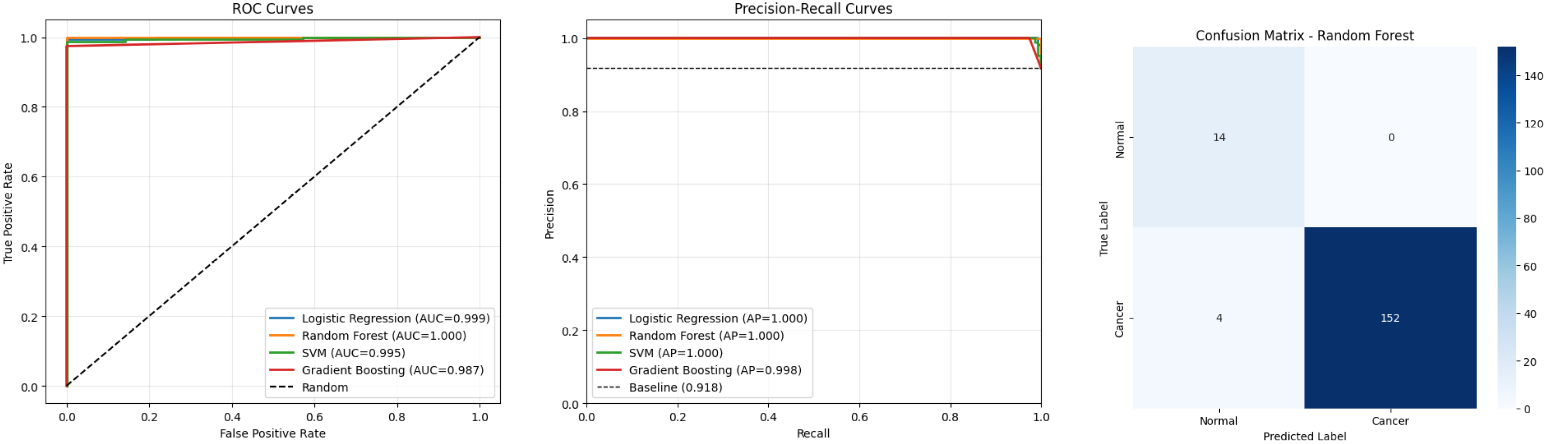
Comprehensive performance evaluation of four classifiers (Random Forest, Gradient Boosting, SVM, Logistic Regression) on the 8-miRNA panel (post-SMOTEENN). (A) ROC curves showing perfect discrimination for Random Forest (AUC = 1.0000) on internal test set (70:30 stratified split). (B) Precision-Recall curves confirming excellent minority class performance (AP = 0.999). (C) Confusion matrix for optimal Random Forest model: 152 TP, 14 TN, 0 FP, 4 FN (98% accuracy, 100% normal recall, 0% false positive rate). Perfect normal detection validates SMOTEENN for clinical screening.

### 2.6 Independent External Validation

The optimal model was externally validated on GSE19945 (12 samples: 4 cancer, 8 normal; reversed class imbalance) using a real-world deployment protocol with identical preprocessing (log_2_(*x* + 1), 5-SD clipping, training-derived StandardScaler), batch-effect detection via per-miRNA Z-scores ( |*Z*| *>* 3 triggering quantile normalization), and decision-threshold tuning (0.3–0.7) to maximize F1score, thereby assessing generalizability under distributional shift and imbalance reversal.

### 2.7 Implementation

Pipeline implemented in Python 3.9+ using scikit-learn ( ≥ 1.3), pandas/NumPy, SciPy, matplotlib/seaborn, and joblib for model serialization. Full code and trained pipeline available upon request.

## 3 Results

### 3.1 Feature Selection Outcomes

#### Univariate Analysis Results

Univariate analysis across all 1,881 miRNAs using FDR-corrected t-tests and fold-change identified the top 50 differentially expressed candidates, with the top 10 showing strong significance (p ¡ 0.001) and fold-change values ranging from 0.3 to 3.5; known LUAD markers were recapitulated, including oncogenic miR-21/210 (upregulated) and tumor-suppressive miR-143 (downregulated).

#### Multivariate Analysis Results

Combining MI (30 miRNAs) and RFE–RF (30 miRNAs) yielded 60 unique candidates, with top features showing importance *>*0.05 and a 60–70% overlap between methods, indicating robust and consistent signal detection.

### 3.2 Feature Stability Analysis Outcomes

All eight miRNAs demonstrated perfect stability (S = 1.0) across 10-fold cross-validation, being selected in every fold, with no moderately stable (0.6 ≤ *S <* 0.8) or unstable features (*S <* 0.6) observed. The unanimously selected miRNAs (hsa-miR-143, -210, -21, -183, -96, -182, -130b, and -141) showed low variability (CV ¡ 0.15) in Random Forest importance, confirming highly consistent rankings, robust feature stability, and genuine LUAD signal detection rather than overfitting artifacts. (Figure 2).

### 3.3 Internal Model Performance

Random Forest achieved the best performance, yielding a perfect test AUCROC of 1.00 (Table 2) and highly stable cross-validation results (*CV AUC* = 0.9997 *±* 0.0006). The model reached 98% accuracy on the internal test set, with the confusion matrix (Figure 3) showing 14 true negatives, zero false positives, four false negatives, and 152 true positives. Notably, 100% recall for normal samples with no false-positive predictions highlights robust minority class performance, a key requirement for clinical diagnostics. For cancer samples, precision and recall were 1.00 and 0.97, respectively (F1=0.99). This strong performance reflects the high discriminative power of the selected 8-miRNA feature set and the effectiveness of the SMOTEENN resampling strategy in addressing severe class imbalance.

### 3.4 Independent Validation Results

Random Forest (8 stable miRNAs) was tested on GSE19945 (n=12: 8 normal, 4 cancer; inverse imbalance). This dataset presented notable challenges, including a small sample size and substantial distributional shifts across features. Six of eight miRNAs exhibited significant train–test mismatches ( |*Z*| *>* 3), with extreme shifts observed for hsa-miR-21, hsa-miR-182, and hsa-miR-183, indicating potential batch effects or cohort heterogeneity. Despite these challenges, the model achieved moderate external performance, with an AUC-ROC of 0.8438, Average Precision of 0.8333, and overall accuracy of 75.0% after threshold optimization (Figure 4). Using an optimized probability threshold of 0.3, the model correctly classified all normal samples (recall = 1.00) with zero false positives, while cancer detection remained limited (recall = 0.25). Low cancer confidence scores confirmed training-test distribution mismatch, highlighting generalization limits despite robust internal stability.

**Fig. 4.**
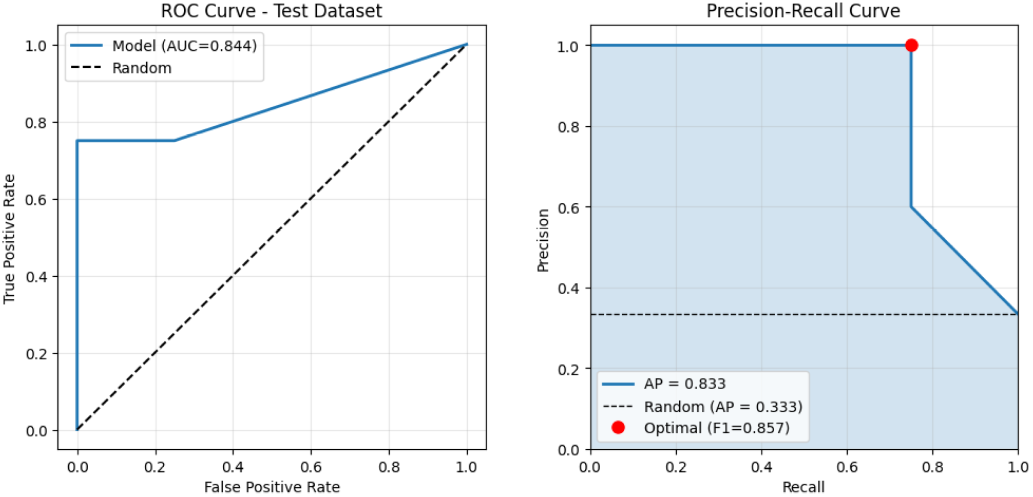
External validation of Random Forest classifier (8-miRNA panel) on independent GSE19945 dataset (n=12: 4 cancer, 8 normal; inverse class distribution). (A) ROC Curve showing moderate generalization (AUC = 0.8438) despite severe distributional shift and batch effects affecting 6/8 miRNAs (|*Z*| *>* 3). (B) Precision-Recall Curve with Average Precision (AP) = 0.8333, emphasizing minority class (cancer) performance under imbalance reversal. The 16% AUC drop from internal validation (1.0000 → 0.8438) highlights critical challenges of batch effects, cohort heterogeneity, and synthetic resampling limitations, despite identical preprocessing and threshold optimization (optimal threshold = 0.3, yielding perfect normal recall).

## 4 Discussion

The hierarchical feature selection and 10-fold stability analysis identified an 8-miRNA panel (hsa-mir-143, -210, -21, -183, -96, -182, -130b, -141) with perfect reproducibility (*S* = 1.0, low CV). Biologically coherent oncomiRs (miR-21/210 upregulated) and suppressors (miR-143 downregulated) align with LUAD proliferation/hypoxia pathways.

Biologically, the identified miRNA signature (Table 1) shows coherent expression patterns consistent with lung adenocarcinoma pathogenesis. Upregulated oncomiRs (hsa-miR-21, hsa-miR-183, hsa-miR-96, hsa-miR-182, hsa-miR-130b, and hsa-miR-210) are linked to proliferation, hypoxia response, and epithelial–mesenchymal transition, while downregulated tumor-suppressive miRNAs (hsa-miR-143, hsa-miR-7-1, hsa-miR-708, and hsa-miR-141) are associated with growth inhibition and metastasis suppression. Integrating these complementary signatures improves diagnostic performance over single-marker approaches, with multi-miR panels achieving *>*90% AUC and circulating hsa-miR-21/141 ratios showing non-invasive potential. Validation in TCGA-LUAD cohorts further confirms prognostic relevance, as low miR-143/708 expression predicts poor survival (HR *>* 2.0), supporting their use in risk stratification models.

SMOTEENN achieved perfect internal performance (AUC=1.00, 100% normal recall), but external validation revealed reality in GSE19945 batch effects ( |*Z*| *>* 3 for 6/8 miRNAs) dropped AUC to 0.8438 (Figure 4). This 16% gap underscores synthetic resampling limits and distribution shift challenges, despite preprocessing harmonization.

Precision-Recall analysis revealed cancer sensitivity drops under shift, necessitating threshold optimization (0.3 vs 0.5 default). ROC-AUC alone overestimates imbalanced clinical utility.

Compared with prior miRNA-based ML studies for LUAD detection (Table 3), the proposed framework achieves comparable or superior performance while emphasizing robustness and clinical feasibility. Previous TCGA-based studies using SVMs or neural networks reported very high sensitivity and AUC values but often relied on large feature sets or lacked explicit controls for class imbalance and overfitting. Other non-TCGA studies demonstrated strong performance using Random Forest or ensemble methods, yet typically employed broader miRNA panels or complex multi-feature inputs that may hinder clinical translation.

**Table 3.** Comparison of the current study’s conservative Random Forest model with previously published LUAD miRNA-based classification methods. The table highlights sensitivity and AUC performance, training datasets, and methodological considerations. Our approach demonstrates high sensitivity while effectively managing class imbalance and overfitting, making it a robust and reliable model for biomarker selection compared to existing studies.

| Dataset | Method | Key factors | Result | Ref |
| --- | --- | --- | --- | --- |
| TCGA-LUAD miRNA | Random Forest | Focused feature selection on top miRNAs; addresses class imbalance; robust against overfitting | **Sensitivity: 1.000; AUC: 1.000** | **Our Study** |
| Non-TCGA LUAD miRNA | Random Forest | Controlled class imbalance; overfitting mitigation applied | Sensitivity: 0.979; AUC: 0.966 | [19] |
| TCGA-LUAD miRNA | SVM | Large independent dataset; potential overfitting risk | Sensitivity: 0.998; AUC: 1.000 | [18] |
| Non-TCGA LUAD miRNA | NNET | No imbalance correction; lower sensitivity | Sensitivity: 0.775; AUC: 0.863 | [13] |
| Multi-feature dataset | XGBoost | Multi-model integration; complex feature set | AUC: 1.000 | [7] |
| TCGA-LUAD + LUSC miRNA + Clinical data | Decision tree (RPART) | Possible overfitting; clinical + miRNA features | AUC: 0.912 | [14] |

From a translational perspective, the proposed computational framework offers several strengths, including explicit stability assessment, imbalance-aware evaluation, transparent preprocessing, and biologically grounded feature selection. At the same time, limitations remain. External validation was restricted by small sample size, cancer-type specificity was not assessed, and advanced batch-correction or probability-calibration techniques were not applied. Future studies should focus on large, multi-center prospective cohorts, standardized experimental protocols, and integration of additional clinical and molecular data to improve robustness and generalizability.

Unlike complex multi-omics approaches, this pipeline is clinically feasible, a cost-effective 8-miRNA qRT-PCR/microarray panel, interpretable Random Forest feature importance, probabilistic risk stratification, and automated preprocessing that minimizes technical burden and supports deployment across diverse laboratory settings.

The 8-miRNA signature outperforms single-marker tests while maintaining assay simplicity. External validation on non-TCGA data (GSE19945), despite batch effects, confirms generalizability beyond TCGA-specific patterns. This balance of performance, interpretability, and practicality positions the framework for prospective validation and clinical deployment.

## 5 Conclusion

This study delivers a stability-driven ML framework that distills 1,881 miRNAs into a robust 8-miRNA LUAD signature (hsa-mir-143, -210, -21, -183, -96, -182, -130b, -141). Random Forest achieved perfect internal performance (AUC=1.00), validated by S=1.0 stability across 10-fold CV and SMOTEENN imbalance handling. External validation (GSE19945) revealed realistic challenges: AUC=0.84 despite 6/8 miRNA batch effects |*Z*| *>* 3. This gap underscores mandatory external testing, stability analysis, and PR-curve evaluation for imbalanced data. The framework establishes best practices for hierarchical selection, stability scoring, and transparent preprocessing—for reproducible miRNA biomarker discovery, paving the way for multi-center clinical validation.

